# Assessing Pathogens for Natural versus Laboratory Origins Using Genomic Data and Machine Learning

**DOI:** 10.1101/079541

**Authors:** Tonia Korves, Christopher Garay, Heather A. Carleton, Ashley Sabol, Eija Trees, Matthew W. Peterson

**Author notes:** Corresponding author (TK).

## Abstract

Pathogen genomic data is increasingly important in investigations of infectious disease outbreaks. The objective of this study is to develop methods for using large-scale genomic data to determine the type of the environment an outbreak pathogen came from. Specifically, this study focuses on assessing whether an outbreak strain came from a natural environment or experienced substantial laboratory culturing. The approach uses phylogenetic analyses and machine learning to identify DNA changes that are characteristic of laboratory culturing. The analysis methods include parallelized sequence read alignment, variant identification, phylogenetic tree construction, ancestral state reconstruction, semi-supervised classification, and random forests. These methods were applied to 902 *Salmonella enterica* serovar Typhimurium genomes from the NCBI Sequence Read Archive database. The analyses identified candidate signatures of laboratory culturing that are highly consistent with genes identified in published laboratory passage studies. In particular, the analysis identified mutations in *rpoS*, *hfq*, *rfb* genes, *acrB*, and *rbsR* as strong signatures of laboratory culturing. In leave-one-out cross-validation, the classifier had an area under the receiver operating characteristic (ROC) curve of 0.89 for strains from two laboratory reference sets collected in the 1940’s and 1980’s. The classifier was also used to assess laboratory culturing in foodborne and laboratory acquired outbreak strains closely related to laboratory reference strain serovar Typhimurium 14028. The classifier detected some evidence of laboratory culturing on the phylogeny branch leading to this clade, suggesting all of these strains may have a common ancestor that experienced laboratory culturing. Together, these results suggest that phylogenetic analysis and machine learning could be used to assess whether pathogens collected from patients are naturally occurring or have been extensively cultured in laboratories. The data analysis methods can be applied to any bacterial pathogen species, and could be adapted to assess viral pathogens and other types of source environments.

## Introduction

Genome sequencing plays an increasingly important role in identifying the origins of disease outbreaks. Disease strain origins are often determined by assessing the genetic relatedness to other strains via phylogenetic analysis or shared genetic markers, and by inferring that closely related strains originate in a common source (1–5). DNA data could also potentially be used to identify the type of environment a strain came from based on adaptive DNA changes. Some environments impose strong selective pressures that tend to cause adaptive DNA changes in certain genes and pathways (6–10). If genome sequence variations that are characteristic of strains adapting to certain environments can be identified, then these could potentially be used to assess which type of environment a strain came from.

One situation where this could be beneficial is in differentiating outbreaks that arise from natural sources from those that have laboratory origins. Disease outbreaks that are the result of naturally circulating strains, due to laboratory accidents, or potentially deliberate events require different types of investigations and response. However, these scenarios are often hard to differentiate, and initially look the same. For example, in the European *Escherichia coli* O104 outbreak in 2011, accidental microbiology lab infections, and infections from deliberate salad bar contamination in Oregon in 1984, the earliest indicator in each event was a sick patient (11–13). It would be advantageous to identify whether an infection was caused by a laboratory strain at this early stage, by analyzing bacterial DNA samples taken from infected patients for evidence of laboratory culturing.

There is substantial experimental evidence for similar DNA changes occurring repeatedly in laboratory culture and in other environments in pathogens, which could potentially be used as indicators of the environment the strains came from. Multiple studies have investigated the evolution of bacteria in laboratory conditions by sequencing DNA from strains before and after passaging in laboratory culture. These studies reveal that some DNA changes are characteristic of adaptation to laboratory culture, both in bacterial species (6,14–16) and in influenza (17–19). The best known of these in bacteria are mutations in the gene *rpoS*, which have been observed in many studies in *E. coli* and in *Salmonella* (16,20–22). In addition, recent studies have found mutations in certain genes of *Burkholderia dolosa* (7) and *Pseudomonas aeruginosa* (23) that are associated with adaptation to patients.

The combination of phylogenetic analysis and large scale genomic data presents an opportunity to discover DNA changes characteristic of certain environments. By determining where on phylogenies certain mutations arise, and how this correlates with environments experienced on those branches on phylogenies, studies can identify parallel DNA changes that are characteristic of certain adaptive pressures. This convergence-based phylogenetic approach has been used to find mutations characteristic of influenza culturing methods (17), adaptive mutations in *Burkholderia* in cystic fibrosis patients (7), and drug resistance mutations in *Mycobacterium tuberculosis* (24). In addition, recent studies have used genomic data from hundreds of pathogen strains to identify DNA polymorphisms affecting antibiotic resistance and virulence, and to predict these phenotypes with machine learning (25,26).

In this study, we investigate whether phylogenetic and machine learning methods can identify genomic signatures of laboratory culturing using publicly available genomic data. We test this approach on 902 genomes of *Salmonella enterica* serovar Typhimurium, a common foodborne pathogen. Our results show that these methods detect signatures of laboratory culturing that are highly consistent with published laboratory passage experiments. Furthermore, a classifier built with these methods can identify a large portion of strains that have experienced substantial laboratory culturing. Finally, we show how these methods can be applied to assessing outbreak strains for laboratory culturing history, and present some evidence suggesting that a set of closely related *Salmonella* outbreak strains may be descended from a laboratory strain.

## Methods & Materials

### Approach for Identifying DNA Signatures of Laboratory Culturing

Our approach is to identify genomic signatures of laboratory culturing based on mutational patterns across a phylogenetic tree (Fig 1). The first step is to recognize which branches of the phylogenetic tree are associated with time in natural conditions and which are associated with time in laboratory culture. If all strains were collected from natural sources, passaged in a laboratory, and subsequently sequenced, then the common ancestors of the strains originated in natural conditions. Consequently, all DNA changes that fall on internal branches of the phylogenetic tree arose in natural conditions. In contrast, DNA changes that fall on terminal branches of the phylogeny arose either in natural conditions (prior to the strain’s collection) or during laboratory passages (after the strain’s collection). Therefore, we expect that genome variants that fall disproportionately on terminal branches of the phylogeny are candidate signatures of laboratory culturing. Our approach is to identify genes, and sets of genes from the same operon, that have excessive mutations on terminal branches of the phylogeny compared to internal branches as candidate signatures of laboratory culturing.

**Fig 1.**
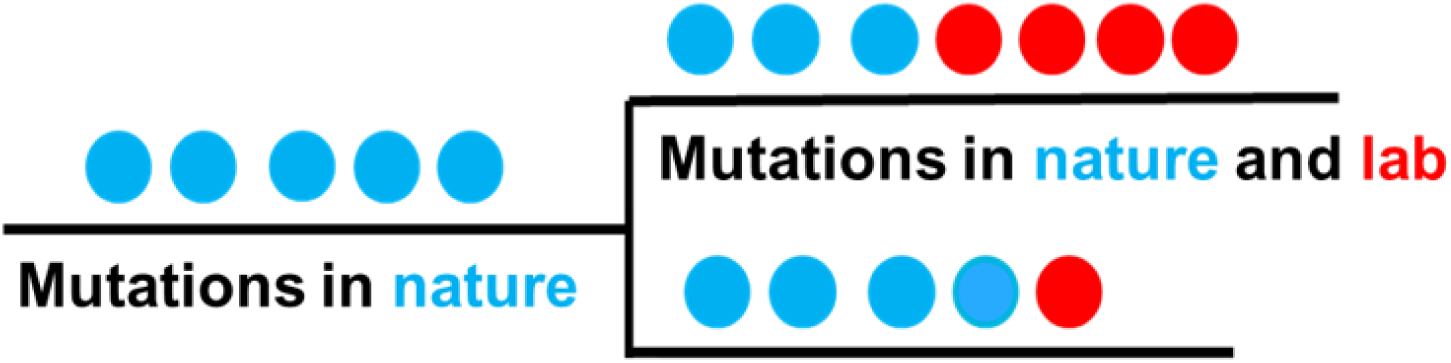
Notional phylogeny branches with mutations that occur in nature (blue) and in laboratory culturing (red).

Our approach includes the following steps:

1. Identify and download sequence read data, and align to a reference strain
2. Identify single nucleotide polymorphisms (SNPs) and deletions
3. Build a phylogeny using the SNP data
4. Map polymorphisms onto the phylogeny. First, reconstruct the ancestral states at the phylogeny nodes. Then map each of the SNPs and deletions onto one or more branches of the phylogeny where the change was most likely to have occurred.
5. Identify candidate signature genes. First, identify the genes that have more mutations on terminal branches, particularly for extensively cultured strains, than on internal branches. Then use machine learning to identify which of these genes, and sets of genes from the same operon, are useful in classifying branches as terminal vs. internal. Genes that contribute significantly to this classification are candidate signature genes of laboratory culturing.
6. Build and test a classification algorithm using the selected genes and gene sets. S1 Fig depicts an overview of the data analysis pipeline and software used to carry out these steps.

### Genome and Strain Data

*Salmonella enterica* serovar Typhimurium was chosen to test this approach because there are a large number of serovar Typhimurium genomes publically available, and for many of these strains we were able to obtain some information about laboratory culturing history. Importantly, serovar Typhimurium has been involved in both naturally occurring and laboratory acquired outbreaks (2,11,13,27–29). In order to facilitate analysis, we selected 948 samples that were associated with paired-end Illumina sequence read data in the NCBI Sequence Read Archive (SRA) and met read depth criteria (see Assembly methods section). The SRA identifiers for the genomes used are listed in S2 Table. Genomes included those generated by public health labs in North America and Europe and genomes which have been published in previous studies (2,27,30–32).

Strains were assigned to culture collection sets in order to group strains that were likely to have experienced similar laboratory culturing histories. These culture collection sets were identified based on strain names, strain collection dates, and the organizations that passaged and housed the strains. We obtained this information through literature searches, from the NCBI BioSample database, and by contacting laboratories that maintained cultures and performed sequencing. Strain collection assignments are given in S2 Table. Information about the methods and the extent of laboratory culturing were obtained by contacting groups that sequenced and maintained the cultures and from publications (30,33,34); this information is given in S3 Table.

### DNA Sequence Read Mapping and Genome Assembly

Raw sequence read data was downloaded in sra format from NCBI SRA (35). Using the SRA toolkit’s fastq-dump (version 2.3.5), sequence reads were extracted to fastq files. Reads were aligned to a reference genome, *Salmonella enterica* serovar Typhimurium LT2 (NCBI reference sequence NC_003197) (36), with the Burrows-Wheeler Aligner (BWA) version 0.7.10 using the aln command (37). See S4 Table for the complete set of parameters used for alignment. In order to ensure that only high-quality samples were used for downstream analyses, we utilized only samples with at least 75% of reads mapped, with at least 90% of the genome covered by reads, and with at least 20x mean read coverage per base.

### Single Nucleotide Polymorphism (SNP) and Deletion Calling

Calls of single nucleotide polymorphisms (SNPs) were performed with the variant calling algorithms in SAMtools version 0.1.19 (38–40). Aligned reads generated with BWA were ordered by genome position and indexed with SAMTools sort and index; pileups and variant calls were generated using mpileup. Any variants with a genotype call Phred-scaled quality of less than 20 were removed unless that variant was present in another sample where it met this quality threshold. All variant calls falling within known phage regions and duplicated genes were removed by filtering with BedTools (41). For each heterogeneous genotype call made by SAMTools, we quantified the number of reads with the reference allele, and the number of reads with the alternate allele. The alternate allele was called in cases where the number of reads with alternate bases was greater than two times the number of reads with reference bases. Otherwise, heterogeneous calls were eliminated from further analysis.

In order to identify deletions, we utilized the Pindel algorithm (42). Pindel identifies paired end reads with one unmapped read and attempts to identify breakpoints spanned by those unmapped reads in order to identify structural variants. We kept deletions identified by Pindel that were supported by 20 or more reads. During visual inspection of regions identified by Pindel, we found that Pindel sometimes identified regions of relatively high coverage as potential deletions. Because of this, we kept only those deletions with a coverage of 10% or less than the mean coverage across the whole genome. Identical deletions in different samples were kept as long as at least one sample contained that deletion such that it met both read support and coverage thresholds. Any variants identified by mpileup that were within the remaining deletions regions were removed from downstream analyses. Only deletions that impacted a single coding gene and/or a single small RNA were included in analyses.

### Phylogeny Construction

After the identification of SNPs in each sample, a SNP matrix was generated and used to produce a FASTA file for each position with a variant in any sample. RAxML was used to generate a maximum-likelihood phylogenetic tree using the standard settings and the GTRCAT generalized time reversible model (43). The tree with the highest likelihood of 20 replicate trees was chosen for further analysis. Strains were pruned from the tree prior to downstream analyses to eliminate replicate strains sharing the same name, laboratory experiment strains, and strain genomes with very long terminal branches, greater than 500 SNPs. Strains that were closely related to the laboratory strain 14028 were also pruned because they were used in a later assessment. This resulted in a set of 902 genomes, plus an LT2 genome used in phylogeny construction.

### Mapping Variants to the Tree

Ancestral reconstruction of variants was performed using the ACCTRAN method in the R package phangorn version 1.99-12 (44). Variants were mapped to branches when the outer node of the branch had the variant state and the more internal node had the ancestral state. Steps were taken to reduce the possibility of variants being assigned to multiple branches due to shared ancestry and imperfect phylogeny construction, rather than to independent events. These cases are most likely when the same variant has been assigned to branches that are close to each other on the phylogenetic tree. To identify these cases, for all variants that were assigned to two or more branches, we calculated the patristic distance between those branches (distance along the tree) and the number of nodes separating them using custom R scripts. If two branches to which the same variant was assigned were separated by less than 0. 0.0012 patristic distance or fewer than eight nodes, then each of the variant assignments to these branches were eliminated from the dataset. The threshold of 0.0012 patristic distance (about 20 SNPs) was chosen to encompass strains from the same outbreak that are closely related and for which the tree topology might be ambiguous. The additional node threshold was selected to eliminate cases where SNPs may have been missed due to low coverage in the sequence data. In addition, because indel variants were more frequently assigned to multiple branches, which could be due to missed indel identification with Pindel, only indels that were assigned to just one branch were kept in downstream analyses. Only internal branches with at least one variant mapped to them were used in downstream analyses. All terminal branches were used, including those with zero variants mapped to them, to incorporate classification of strains with no unique variants. These analyses were performed using custom R scripts.

### Assigning Mutations to Genes, Grouping Mutations by Operon, and Selection of Features

To identify mutation effects at the gene level, we reduced the variant set to a maximum of one mutation per gene per branch. We utilized SnpEff (45) to predict which genes each of the identified genetic variants affected. Any deletion or nucleotide polymorphism, whether synonymous or non-synonymous, was considered to have an effect. Synonymous changes were included because of evidence that synonymous changes can impact mRNA stability and fitness (46). We also identified mutations in small RNAs using the positions of small RNAs listed in Sittka et al., Table S3 (47). Because we seek patterns associated with laboratory culturing and not environments that occur in nature, we used only genes that were rarely affected on internal branches and had substantially more mutations on terminal branches than on internal branches. Specifically, we included genes that had mutations assigned to no more than four internal branches, had at least three mutations assigned to terminal branches, and had more than four times as many mutations assigned to terminal branches than to the internal branches.

Mutations in genes that closely interact with each other can have similar lab-adaptive phenotype effects, such that a mutation in either gene could constitute a signature. Therefore, we sought a simple way to pool potentially interacting genes to create composite features. Because genes within the same operon are more likely to be involved in similar processes than pairs of genes at random, we assigned genes to operons using ProOpDB (48). We then pooled mutations at the operon level for genes that met the following criteria: gene had mutations assigned to no more than two internal branches, had at least four mutations assigned to terminal branches, and had more than four times as many mutations assigned to terminal branches than to the internal branches. Mutations in genes that did not meet these criteria were not included in the operon features. Only operons that had two or more genes that met these criteria were included as features (operons with one such gene were already covered by the individual gene criteria above.) As was done for individual genes, for operon gene sets we included a maximum of one mutation per operon gene set per branch.

### Analyses of Mutational Patterns in Strain Culture Collections and Selection of Strains for Building a Classifier

Only two of the culture collections were known to have experienced substantial laboratory passage, and information about passage history was unavailable for multiple culture collections in our dataset. Therefore, we sought to identify additional strains that may have experienced substantial laboratory culturing in order to increase the number of samples for model building and identification of signatures. To identify additional strains that are likely to contain laboratory acquired mutations, we performed unsupervised clustering on all of the branches of the phylogeny, and examined assignments to clusters. We first calculated proximities among all branches with unsupervised random forest classification using the randomForest package version 4.6-10 (49) in R version 3.1.3 (50). This was done using the gene and gene set features described above. We then performed k-medoid analysis using the *cluster* package in R (51,52). Each strain was assigned to one of two clusters. We observed the cluster assignment patterns for terminal branches from the two old reference collections and for internal branches, and assessed the other culture collections for their similarity to each of these patterns.

The strains used as positive examples in analyses for identifying candidate signature genes and classifier building met one of three criteria: 1) The strain belonged to one of the two laboratory reference collections dating back to the 1940’s or 1980’s. 2) The strain belonged to a culture collection that had a high percentage of its strains assigned to the cluster representative of the two reference culture collections and there were more than ten strains in the collection in our dataset. 3) The strain was assigned to the cluster representative of the two reference culture collections and was not from one of the culture collections reported to be passaged less than seven times and stored frozen.

### Identification of Candidate Signatures

Candidate signatures were identified by using the R *Boruta* package version 4.0.0, which identifies features that significantly contribute to random forest classification (50). The standard Boruta settings were used, including p-value <0.01 for confirmation of features. The algorithm was used to classify internal branches versus terminal branches for the strains selected based on the unsupervised cluster analysis. The model included both individual genes and sets of genes from operons as features, which were selected using the criteria described above. For all genes and gene sets that were not rejected in any of the five Boruta runs, variable importance scores (mean decrease in accuracy) were calculated. This was done by including all of the non-rejected genes and gene sets as features in a random forest model and calculating importance using the random forest package.

### Evaluation of Candidates by Comparison to Mutations Observed in Laboratory Experiments

To compare candidate DNA signatures from our analyses to DNA changes observed in laboratory evolution experiments, data were assembled on genes that mutated in published laboratory passaging experiments (6,14,15,53–66). This included genes that mutated in any laboratory experiment in *Salmonella enterica*, and genes that were reported to have mutated in at least two independent replicates or studies in *E. coli*, which is closely related to *Salmonella*. A list of these genes is given in S5 Table. For candidate signature genes that were not on the list of genes that mutated in laboratory experiments, we used the STRING database (67) to investigate whether the candidate signature gene interacted with any of the genes identified in laboratory experiments. We used an interaction score of 0.9 as the threshold.

### Building and Testing a Classifier

We built classifiers using random forests with 2000 trees with the R package randomForest (49), using internal branches (negative examples) versus terminal branches from strains selected in the cluster analysis (positive examples). To test the ability of these methods to identify laboratory cultured-strains, we performed a leave-one-out cross-validation (LOOV) analysis using the caret package to create folds (68). The LOOV analysis performed feature selection and classifier building on the training set, and tested on the left out branch. Feature selection used the same criteria as described above. Predictions for the left out branches were compiled to calculate recall and false positive rates. In addition, predictions for the terminal branches for the 1940’s and 1980’s reference set strains and for the internal branches were used to build a receiver operating characteristic (ROC) curve using the *AUC* R package (69).

### Using the Classifier to Assess Outbreak Strains for Laboratory versus Natural Origins

We applied the classifier to outbreak strains closely related to serovar Typhimurium laboratory reference strain 14028. This set included ten strains which were associated with acquired laboratory infections of *Salmonella* and ten strains associated with a 2009 foodborne outbreak associated with bagged lettuce (S6 Table). All of these strains were indistinguishable from the strain 14028 by pulsed field gel electrophoresis (PFGE) and were sequenced on the Illumina MiSeq (Illumina Inc., San Diego, CA) using 2x250 bp chemistry. Sequence data for these strains is available in NCBI SRA; see S6 Table for the identifiers. We also identified eleven additional genomes in NCBI SRA that were closely related to these strains (S6 Table). For these 31 strains and several related strains used as outgroups, we identified variants using methods described above, except that a higher threshold for calling a SNP was used. In order for a SNP to be called at a location, it had to have a phred-scaled quality score of at least 100 in at least one of the genomes in this set, and calls to no more than one nucleotide variant in the larger set of genomes used to build the classifier. We built a phylogeny using the methods described above, but used SNPs outside of coding genes in addition to SNPs within coding genes to incorporate additional variation. Variants were mapped to the phylogeny as described above. Each branch was then tested using a random forest classifier built from the original dataset of 902 genomes with the methods used in the LOOV analysis, which yielded a prediction value for each test branch. P-values for the test branches were then determined by calculating the fraction of internal branches in the LOOV analyses that had a higher prediction value than the test branch prediction value.

## Results

### *Salmonella* serovar Typhimurium Polymorphisms Mapped to Phylogeny Branches

From 902 serovar Typhimurium genomes, the analysis pipeline identified 17,229 SNPs and 492 deletions in coding genes and small RNAs, of which 17,058 SNPs and 402 deletions were mapped onto a phylogeny (tree in S7 File). Ninety-eight percent of the mapped SNPs were assigned to just one phylogeny branch, and 99.8% were assigned to three or fewer phylogeny branches, which suggests that the ancestral reconstruction and filtering steps resulted in a data set with few ambiguous SNP assignments to branches. In the set mapped to the phylogeny, polymorphisms occurred in 3,456 out of 4,620 annotated protein coding genes in the serovar Typhimurium LT2 reference genome and in 67 small RNA genes. After reducing the mutations to a maximum of one mutation effect per gene per branch assignment, there were 17,177 gene mutation events on branches, which were used for identifying candidate signature genes in further analyses. Sixty-two percent of these were on terminal phylogeny branches and 38% were on internal phylogeny branches.

### Mutation Patterns Consistent with Laboratory Culturing in Strain Culture Collections

To assess whether some strains showed distinctive mutation patterns that could be due to laboratory mutations, we performed unsupervised cluster analysis on all the phylogeny branches. We asked whether terminal branches for some strains clustered separately from internal branches; internal branches in this phylogeny represent mutation patterns under natural conditions. In the two strain collections known to have long laboratory histories, a reference collection originating from the 1940s (LT) and a reference collection originating from the 1980s (SARA), greater than 65% of the strain terminal branches were assigned to cluster 2, while only 1% of internal branches were assigned to this cluster (Fig 2). In contrast, for the six strain collections reported to have been experienced little laboratory culturing (passaged only few times and stored frozen), terminal branch clustering results more closely resembled internal branch patterns. Strain collections with unknown laboratory histories had numbers that ranged from similar to internal branches to numbers similar to the extensively cultured strain collections. These results indicate that strains from collections known to be extensively cultured, as well as from a few collections with unknown lab passage history, exhibit mutational patterns that are consistent with the presence of distinctive, laboratory acquired mutations. All strains from the four culture collections that had at least 40% of their strains assigned to cluster 2, which included the two reference strain collections and the collections N and O, shown in Fig 2, were used in further analyses to identify candidate signature genes. In addition, strains from other collections that were assigned to cluster 2 and not from the six collections reported to have experience little laboratory culturing were also used in downstream analyses as positive examples.

**Fig 2.**
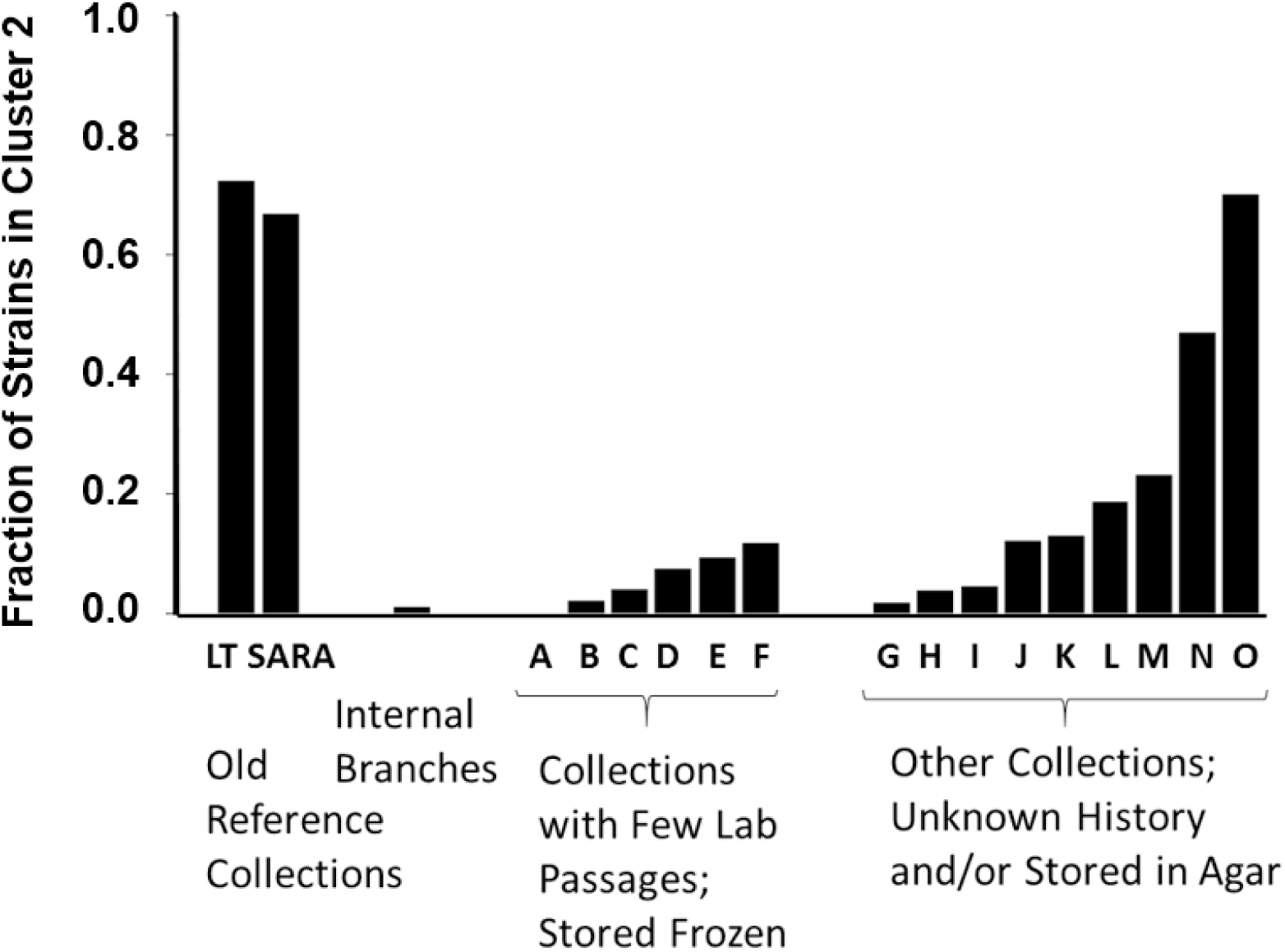
Fraction of branches assigned to one of two clusters for strain collections and internal branches. Except for LT and SARA collections, only collections that contain at least 20 strains are shown. Results are from unsupervised random forest classification and k-medoids clustering.

### Candidate Signatures of Laboratory Culturing

Using a random forest classifier, we identified candidate signature genes and operon gene sets that are highly consistent with results from published laboratory experiments (Fig 3). These genes and operon gene sets were confirmed as significant in all five replicate Boruta, and ranked by the size of the contribution to differentiating internal branches and terminal branches from the strains selected in the unsupervised cluster analysis. The six top-ranked features contained genes that mutated in prior lab studies: *rpoS*, *hfq*, *rfbJ*, *acrB*, and *rbsR*. The genes that made the largest contributions were *rpoS*, a gene well-known to mutate during laboratory passaging, and *hfq*, which is known to interact with *rpoS*. Changes in *rpoS* occurred 31 times on terminal branches of the phylogeny and were never observed on internal branches (S8 Table). Two other genes that interact with *rpoS* were also identified as contributing: *dksA*, which is an RNA polymerase-binding transcription factor, and *nlpD*, the gene that contains the promoter for *rpoS*. In addition, a small RNA that upregulates *rpoS*, *sraH*, was identified as a potential, weaker candidate signature (S8 Table.) Eight of the 34 genes (24%) in the top twenty candidate signatures have been identified in published laboratory studies, which is a far higher proportion than lab study genes in the genome at large (4.4%, 202 genes found in published lab studies out of 4621 annotated genes in serovar Typhimurium LT2; one-sided Fisher exact test, p<0.0001.) In addition, seven other candidate signature features have strong relationships with genes identified in the lab studies (Fig 3). An additional 51 genes and operon gene sets were not rejected as candidates in any of the Boruta runs, and may also include potential candidate signatures (S8 Table). Overall, these results indicate that the bioinformatic analyses of publicly available genomes successfully identified signature genes of laboratory culturing.

**Fig 3.**
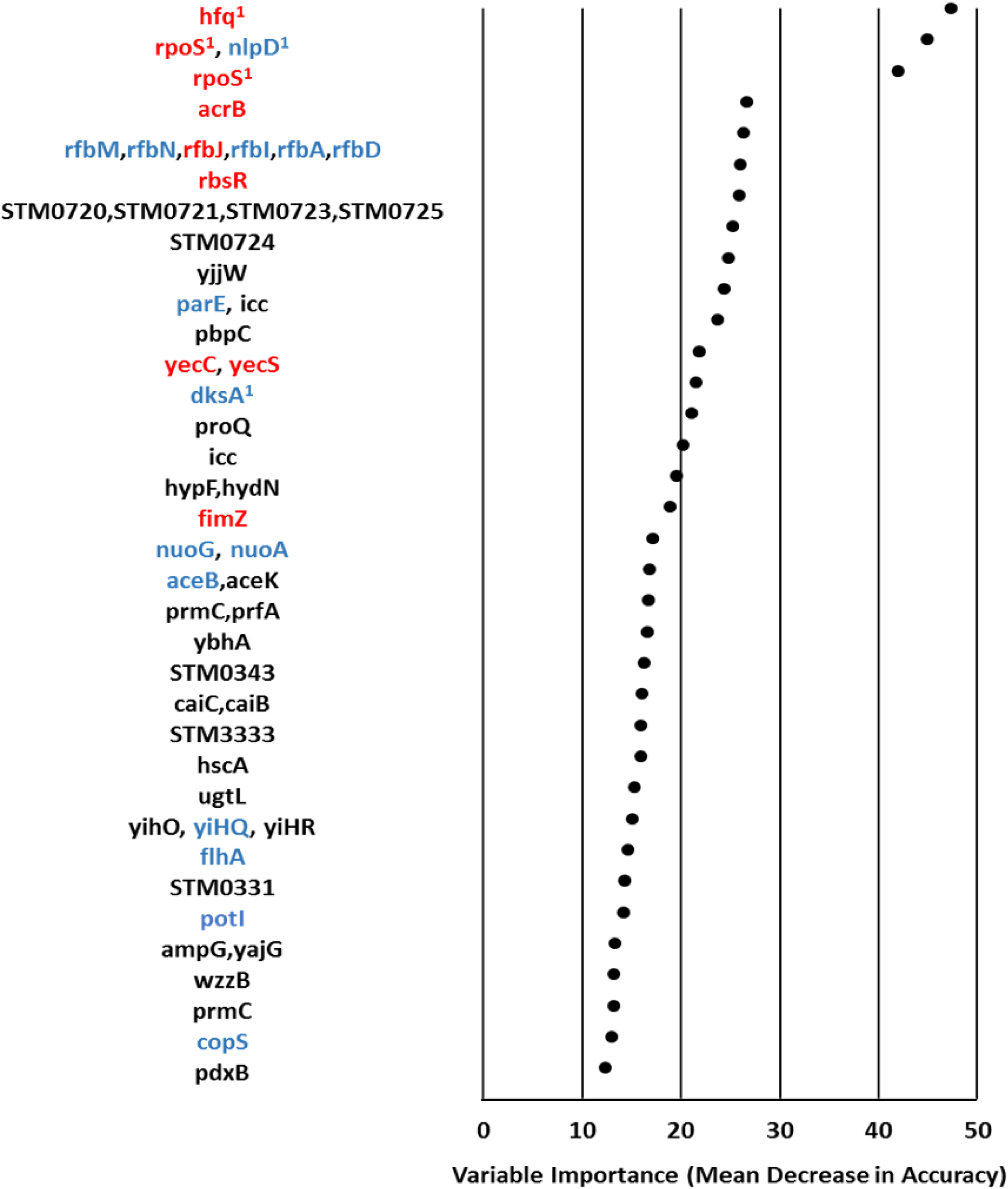
Candidate signature genes and operon gene sets with variable importance scores. Red indicates gene mutated in a published laboratory passage experiment in *Salmonella* or *E. coli*. Blue indicates gene is strongly associated in the STRING database with another gene that mutated in a published laboratory passage experiment. Black indicates no association found with published laboratory study genes. 1 marks *rpoS* and genes that are known to interact with it.

### Performance of the Classifier

We built random forest algorithms to classify strains as having experienced laboratory culturing versus natural origin, and assessed performance using leave-one-out cross-validation. For the culture collection with the most extensive laboratory passaging (LT), the classification algorithms detected half of the strains with a 2 % false positive rate (Table 1), and 78 % of LT strains were detected at a 10% false positive rate (Table 1). The ROC curve in Fig 4 shows results for the two old reference strain collections, and the area under the curve is 0.89. Results for the two old reference collections and the two other collections identified in the unsupervised cluster analysis are in Table 1. The classifier performed better on branches with less than ~50 SNPs long than on longer branches, due to a high number of false positives for long internal branches (S9 Figure). For culture collections reported to have been lab-passaged very little, and other culture collections with unknown laboratory histories, results were similar to internal branches (Table 1).

**Table 1.** Results from Leave-one-out Cross-validation (LOOV).

| Strain Culture Collection | Number<br>of Strains | Percent Predicted to<br>Have Lab Origins at a<br>2% False Positive<br>Rate | Percent Predicted to<br>Have Lab Origins at a<br>10% False Positive<br>Rate |
| --- | --- | --- | --- |
| LT (collected in 1940's) | 18 | 50 | 78 |
| SARA (collected in 1980's) | 15 | 27 | 40 |
| Collection N | 32 | 44 | 63 |
| Collection O | 20 | 35 | 55 |
| Internal Branches (false positive rate) | 452 | 2 | 10 |
| Collections Reported as Minimally Cultured | 235 | 2 | 6 |
| Other Collections with Unknown Histories | 582 | 4 | 10 |

**Fig 4.**
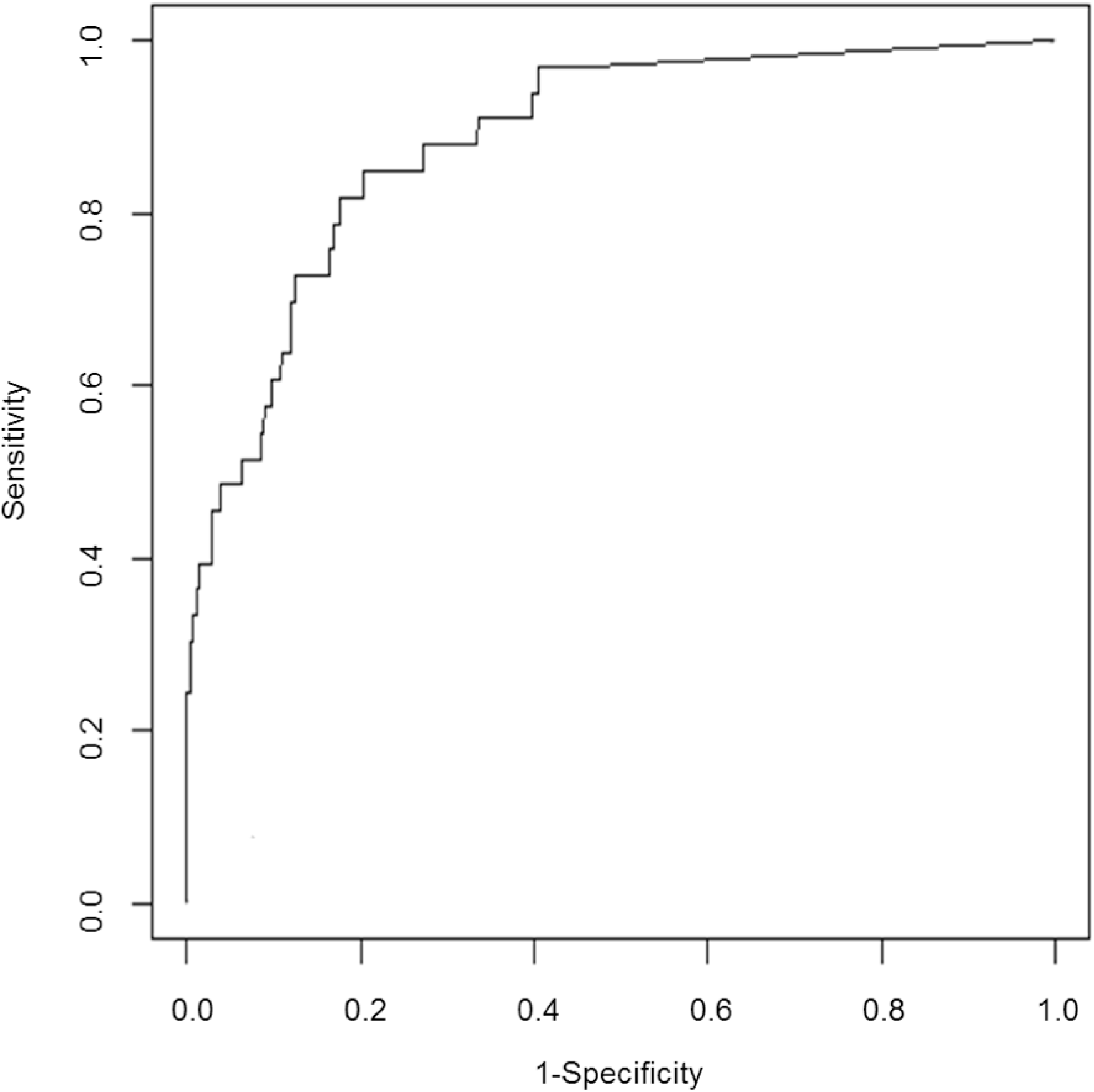
ROC curve showing results for strains from the two old reference strain collections. Strains from LT and SARA collections are treated as true cases and internal branches as negative cases. Results are from the LOOV analysis.

Among these culture collections with at least twenty strains, results ranged from zero to 19 % being classified as lab-origin at a 10% false positive rate. Overall, these results indicate that a classifier can identify a substantial portion of strains from some culture collections that have experienced extensive laboratory culturing, and identifies few strains from culture collections with more minimal laboratory culturing.

### Assessment of Laboratory Culturing in *Salmonella* serovar Typhimurium Strains Closely Related to Laboratory Strain 14028

We assessed evidence of laboratory culturing in strains closely related to laboratory stock strain serovar Typhimurium 14028 by constructing a phylogeny of these strains and applying the classifier built on the other set of strains. Phylogenetic analysis revealed that all of the 14028 related strains were very closely related to each other, with little phylogenetic structure among them (Fig 5). Notably, the strains associated with acquired laboratory infections were interspersed in the phylogeny with those associated with a 2009 foodborne outbreak. In total, 35 mutation events, including 31 SNPs and four deletions, were mapped to the branches within this clade. The number of unique variants per strain ranged from zero to eight. The small amount of variation among the strains in this clade suggests that all of these strains are descended from a recent common ancestor.

**Fig 5.**
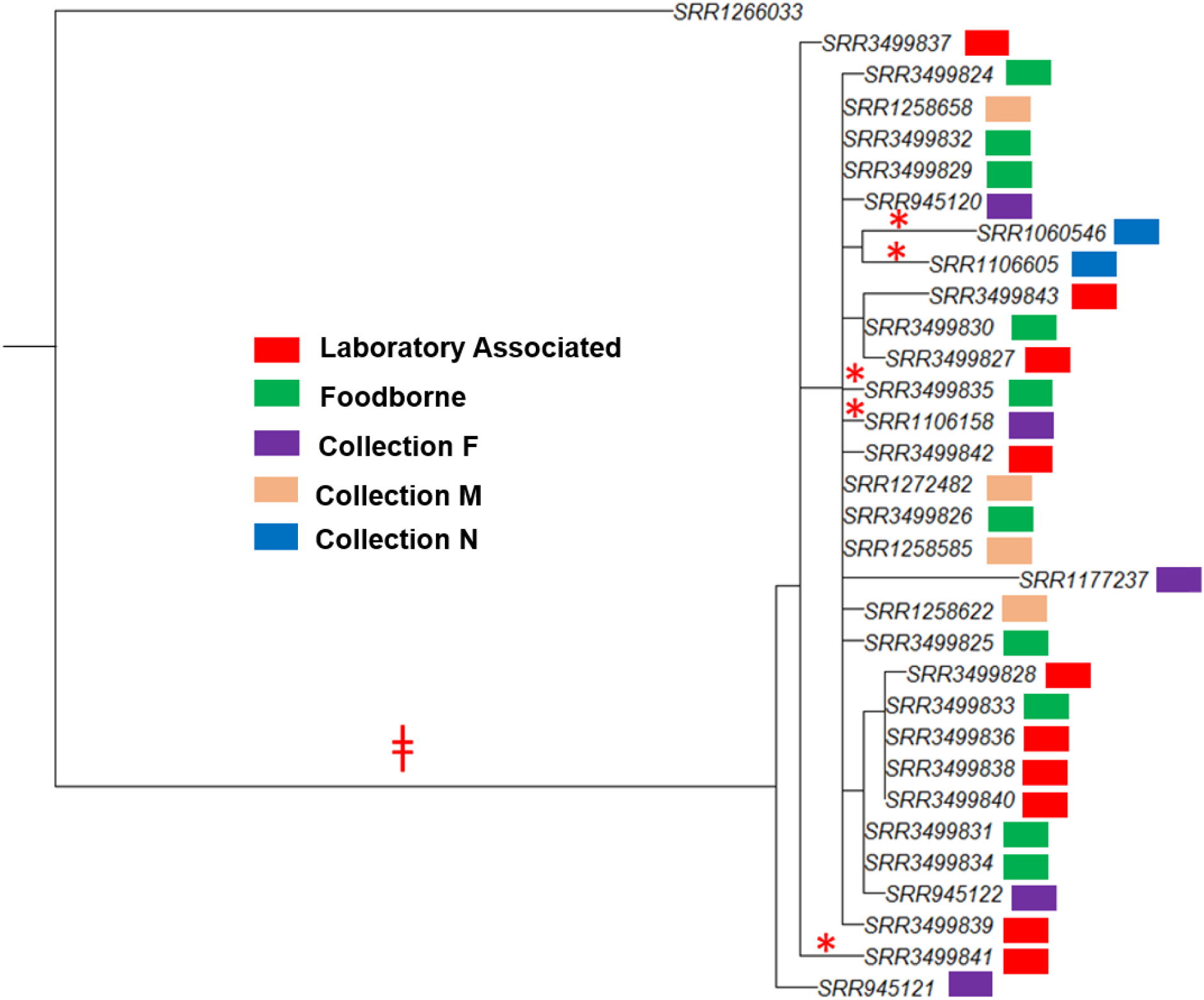
Phylogeny of serovar Typhimurium strains closely related to strain 14028 with results from the classifier.

The classifier detected some evidence of laboratory culturing on the internal branch leading to this clade (Fig 5), suggesting that the common ancestor of this clade may have experienced laboratory culturing. The prediction value generated by the algorithm corresponded to a false positive rate of 6.7%. The 20 SNP mutations mapped to this branch included a mutation in the gene STM0725, a putative glycotransferase that is part of a candidate signature operon gene set, and *pdx*B, another candidate signature gene (Fig 3).

The classifier also detected evidence of laboratory culturing in five individual strains within this clade (Fig 5). All five strains had mutations in the highly ranked, interacting candidate signature genes *rpo*S, *nlpD*, and/or *hfq*. Two of these strains came from strain culture collection N, which exhibited strong evidence of laboratory culturing in the larger set of strains (Fig 2 and Table 1); therefore, these mutations may reflect passaging after strain isolation. The three other mutations occurred in one strain associated with a laboratory acquired infection, one strain associated with a community college microbiology class (SRR1106158, personal communication, A. Perez Osorio, Washington State Department of Health) and one strain associated with a foodborne outbreak.

Symbols mark branches where the classifier detected some evidence of laboratory culturing. *, false positive rate of less than 3%. ǂ, false positive rate of less than 10%.

## Discussion

The combination of large-scale DNA data and machine learning has recently been used to identify signatures of antibiotic resistance and predict virulence in pathogens (25,26). This study describes a way in which genomics and machine learning can also be used for insight into the origins of disease outbreaks. We present analysis methods that identify signatures of laboratory culturing by identifying parallel evolutionary changes in large-scale, publicly available genome sequence data. We show that these genetic signatures can be used to assess whether pathogens have experienced substantial lab culturing. While our analysis was performed on *Salmonella* genomes, our approach is generalizable and can be used for analyzing the origins of other pathogens.

One potential use of these methods is in the investigation of outbreaks where laboratory acquired or deliberate infections may be suspected. In cases where there is circumstantial information suggesting that an outbreak may not be natural, these methods could be used to evaluate whether a pathogen collected from a patient shows signs of having come from a laboratory. This could indicate whether an investigation of the outbreak as a potential laboratory exposure or other laboratory-origin event is warranted. Given that outbreaks of laboratory-origin are very rare, the classifier would have a low positive predictive value when applied to outbreaks at large and consequently this method would probably not be effective for general screening of all outbreak pathogens without large increases in classifier precision. Another potential application of these methods is in identifying laboratory-acquired mutations in culture collections, in order to account for these in vaccine and drug development and in other scientific investigations.

When we applied these methods to a set of *Salmonella* strains closely related to laboratory strain 14028, the classifier results detected some evidence of laboratory culturing in the ancestral strain of this set. Together, these classifier results, combined with the presence of known laboratory strains in this clade and the low variation within this clade, suggest that all of the strains in this clade may be descended from a laboratory strain. The serovar Typhimurium strain 14028 was originally collected in 1960 and has been a laboratory stock strain for many decades (63). It has been used as a reference strain in university laboratory classes and in diagnostics, has been associated with laboratory acquired infections, and was even used in a deliberate contamination of salad bars in 1984 (11,13,28). Consequently, if these strains are all derived from the original laboratory strain, they may reflect multiple laboratory escape events over time.

Comparisons to published laboratory passaging experiments in *E. coli* and *Salmonella* show that our method identifies genetic signatures of laboratory culturing. In particular, *rpoS,* and genes known to interact with it, were the strongest signatures in our set. This is consistent with many lab studies that have identified mutations in *rpoS*, and, to a lesser extent in *hfq*, that occur during lab culture (55,58,60,62–64). Our study expands this set to include the genes *nlpD*, which contains the *rpoS* promoter, and *dksA*, which interacts with both *rpoS* and *hfq*. We also identified mutations in *acrB* as a strong signature of laboratory culturing, which is consistent with recently observed laboratory mutations in *acrB* and its interacting gene *acrA* (58). Other genes found in laboratory culturing experiments that made substantial contributions to classification are *rfb* genes and *rbsR* (15,54,61,63). Several genes not found in laboratory studies were also identified as strong candidate signatures, including a set of five putative glycosyl transferase genes from a single operon. The candidate signatures identified in this study would benefit from further experimental validation.

It is likely that there are more genes characteristic of laboratory culturing that we were unable to detect. First, the dataset included a diverse set of culture collections subject to a variety of culture methods, and experimental studies indicate that whether certain genes mutate or not is dependent on growth conditions, such as stationary phase laboratory culturing and stab cultures (16). Types of mutations that occur in growth conditions that were rare in our sample would be unlikely to be detected. Second, experiments indicate that gene mutations in laboratory culture depend heavily on the genetic background of that strain (16). Thus it is likely that there are adaptive characteristic mutations the analysis did not identify because they are specific to certain backgrounds, or a small set of backgrounds such that they do not appear a sufficient number of times in our sample.

The classifier identified many of the strains from extensively cultured collections as having been laboratory passaged, but also identified a much smaller portion of strains from some other collections. Results for strains from culture collections that experienced only isolation culturing steps resembled internal phylogeny branches, suggesting that detection of only a small amount of laboratory culturing might not generally be possible by this method.

There are several extensions that would likely identify additional DNA signatures of laboratory culturing and improve classification. First, our dataset contained only 33 genomes from culture collections known to have experienced substantial lab culturing and an additional 99 selected in the unsupervised cluster analysis. Inclusion of more genomes known to have experienced substantial laboratory culturing would increase the ability to identify genes that mutate less frequently as signatures. Second, our analyses only included DNA segments present in the reference genome chromosome and left out phage sequences. The inclusion of additional DNA segments, such as from plasmids and chromosomal regions present in some strains but not in the LT2 reference, should yield additional features that would also enhance recall and specificity. In addition, use of a different reference strain that is more closely related to currently circulating serovar Typhimurium strains might also yield additional signatures. Finally, our analyses suggest that sets of interacting genes are potential candidate signatures, and feature creation that incorporates mutations at the level of sets of interacting genes, beyond operons, may enhance classification. Overall, a combination of more genetic data and improved feature engineering is likely to improve sensitivity and specificity.

Our analysis also suggests that it may be possible to discover signatures of laboratory culturing and build a classifier even when there is no information available about the laboratory history of strains in the dataset. By performing unsupervised classification on genes and operon gene sets that have mutated more on terminal than on internal branches, analyses can identify strains that show patterns that are distinct from natural patterns for use in building a classifier. This is important because information about laboratory culturing history is rarely captured in publicly available databases, and this enables the use of more extensive data. Nevertheless, test cases and information about mutations in laboratory culture in related strains are important to confirm that the model is identifying laboratory signatures, and not signatures of another environment type.

Our analytical approach can be applied to any pathogen species, and could be adapted for identifying more than just a history of laboratory culture. The data analysis pipeline can be readily applied to other bacterial species and adapted for viral species. The methods could also be modified for classification of other types of environmental sources, such as determining whether a pathogen came from cattle or chicken hosts. For this, source environments would be mapped onto the phylogeny differently than for laboratory culturing, but the other steps would apply. With further development, this approach potentially offers a way to infer the type of environment a pathogen came from, and could be a useful complement to inferences based on DNA relatedness in disease outbreak investigations.

## Acknowledgements

We thank D. Brown, D. Boxrud, J. Lahti, A. Mather, E. De Pinna, K. Sanderson of the Salmonella Genetic Stock Centre, D. Toney, and W. Wolfgang for laboratory culture history information. We also thank M. Colosimo for helpful review.

## Disclaimer

The findings and conclusions in this report are those of the authors and do not necessarily represent the official position of the Centers for Disease Control and Prevention. Use of trade names is for identification only and does not imply endorsement by the Centers for Disease Control and Prevention or by the U.S. Department of Health and Human Services.

## Supporting Information

S1 Figure. Schematic of the data analysis pipeline

S2 Table. SRA identifiers for genomes in the large phylogeny

S3 Table. Laboratory culturing histories of the strain collections S4 Table. Parameters used for BWA sequence alignment

S5 Table. Genes that mutated in *Salmonella* and *E. coli* in published laboratory culturing experiments S6 Table. SRA identifiers for strains closely related to serovar Typhimurium 14028

S7 File. Phylogeny of 902 serovar Typhimurium strains and LT2

S8 Table. Genes and sets of genes identified as potential candidate signatures

S9 Figure. Relationship between branch length and predicted probability of extensively lab culturing

